# Novel antibodies for the simple and efficient enrichment of native O-GlcNAc modified peptides

**DOI:** 10.1101/2021.05.28.446228

**Authors:** Rajan A. Burt, Borislav Dejanovic, Hayley J. Peckham, Kimberly A. Lee, Xiang Li, Johain R. Ounadjela, Anjana Rao, Stacy A. Malaker, Steven A. Carr, Samuel A. Myers

## Abstract

Antibodies against post-translational modifications (PTMs) such as lysine acetylation, ubiquitin remnants, or phosphotyrosine have resulted in significant advances in our understanding of the fundamental roles of PTMs in biology. However, the roles of a number of PTMs remain largely unexplored due to the lack of robust enrichment reagents. The addition of N-acetylglucosamine to serine and threonine residues (O-GlcNAc) by the O-GlcNAc transferase (OGT) is a PTM implicated in numerous biological processes and disease states but with limited techniques for its study. Here, we evaluate a new mixture of anti-O-GlcNAc monoclonal antibodies for the immunoprecipitation of native O-GlcNAcylated peptides from cells and tissues. The anti-O-GlcNAc antibodies display good sensitivity and high specificity toward O-GlcNAc-modified peptides, and do not recognize O-GalNAc or GlcNAc in extended glycans. Applying this antibody-based enrichment strategy to synaptosomes from mouse brain tissue samples, we identified over 1,300 unique O-GlcNAc-modified peptides and over 1,000 sites using just a fraction of sample preparation and instrument time required in other landmark investigations of O-GlcNAcylation. Our rapid and robust method greatly simplifies the analysis of O-GlcNAc signaling and will help to elucidate the role of this challenging PTM in health and disease.

## Introduction

O-GlcNAc is the monosaccharide addition of N-acetylglucosamine to serine and threonine residues of nuclear, cytosolic, and mitochondrial proteins. Distinct from N- and O-linked ER and Golgi glycosylation pathways, O-GlcNAc is considered to be a dynamic, regulatory, intracellular post-translational modification (PTM). Unlike other types of glycosylation, O-GlcNAc is not elaborated beyond the initiating modification. Understanding the functional relevance of specific O-GlcNAc sites has proven challenging for several reasons. The sole enzymes responsible for the addition and removal of O-GlcNAc, OGT and OGA/MGEA5, respectively, are essential for embryonic development (1,2). Tissue-specific deletions of either gene present different and often deleterious effects depending on the context (3–7). The mechanisms underlying these phenotypic changes are likely pleiotropic as O-GlcNAc signaling affects nearly every cellular process (Reviewed in (8)). As such, having an experimental strategy to confidently map O-GlcNAc sites across the proteome would be a powerful tool to dissect the functions of this diverse PTM.

Mapping O-GlcNAcylation affords its own challenges including enriching O-GlcNAc-modified peptides from complex biological mixtures and unambiguously identifying sites of modification. Regarding the latter, the glycosidic bond between the GlcNAc and the hydroxyl-containing residue is labile. Typical collisional activation conditions (CID, HCD, etc.) result in near stoichiometric loss of the GlcNAc moiety from the precursor glycopeptide. This removes the information required for site-assignment (9,10). Fragmentation using electron capture dissociation (11), and its more recent replacement electron transfer dissociation (ETD) have been well-documented to preserve the O-glycosidic bond while breaking the peptide backbone to enable unambiguous modification site assignment (12–14). ETD, however, is inherently slower than HCD and is not compatible with widely used isobaric peptide labeling quantification strategies such as iTRAQ and TMT. Consequently, HCD has been used in combination with ETD by using an HCD-product-dependent ETD acquisition method (HCD-pd-ETD), where sugar oxonium ions and fragments thereof are detected in the HCD scan triggering ETD on the same, re-accumulated precursor (9,15).

Several strategies have been developed to enrich O-GlcNAc-modified peptides, though most have notable drawbacks (Reviewed in (16)). Metabolic labeling with monosaccharides bearing bioorthogonal chemical moieties has been used successfully in multiple iterations (17–19). By culturing cells with chemically-modified monosaccharides, these sugars are incorporated into metabolic pathways that eventually modify proteins. Modified peptides or proteins can then be coupled to an affinity handle for subsequent enrichment and analysis by immunoblotting or mass spectrometry. What metabolic labeling gains in sensitivity, it loses in applicability, being incompatible with sample sources that cannot be cultured in the laboratory. Chemo-enzymatic approaches have also been successful, where azide or alkyne-modified N-acetylgalactosamine (GalNAc) is enzymatically transferred by a mutant GalT1 to terminal GlcNAc moieties before being coupled to an affinity handle (14,20). The widespread adoption of this approach, however, has been hindered by its complexity.

Enrichment of native O-GlcNAc-modified peptides is important to expand our knowledge of this PTM’s function, allowing cells and/or tissues from mouse models or human patients to be compared across biological states. Phenylboronic acid can coordinate the cis-diol moiety of glucose for enrichment of native O-GlcNAc-modified peptides, though its sensitivity and specificity have not been fully explored (21,22). Wheat germ agglutinin (WGA), a lectin that has a weak affinity for GlcNAc, has been used successfully for O-GlcNAc-modified peptide enrichment across a wide range of cells and tissues (23–25). Although WGA-based lectin weak affinity chromatography (LWAC) has provided some of the largest datasets of O-GlcNAc-modified peptides (24,26,27) its lack of specificity is well-documented, and the amount of input peptide needed can be prohibitive (26,28,29).

A simple and sensitive strategy to enrich for native O-GlcNAc-modified peptides in cells and tissues would greatly facilitate the study and understanding of O-GlcNAc signaling on a global and local (site-specific) scale. Here, we characterize a newly developed mixture of anti-O-GlcNAc monoclonal antibodies capable of enriching native O-GlcNAc-modified peptides from cells and tissues using peptide input amounts compatible with many sample types. The antibody mixture shows excellent specificity towards O-GlcNAc-modified peptides, reducing the number of co-enriched glycan-containing peptides that are common with other enrichment strategies. We show that these anti-O-GlcNAc antibodies can be coupled with offline fractionation to enable deep characterization of the O-GlcNAc proteome from mouse brain tissue. We believe this commercially available enrichment reagent will be a breakthrough for our ability to map global, site-specific O-GlcNAcylation patterns across a diverse range of biological samples.

## Experimental Procedures

### Anti-O-GlcNAc antibody development

Polyclonal antibodies were produced by immunizing New Zealand White rabbits with randomized peptide libraries containing serine and threonine residues modified with O-linked GlcNAc coupled to KLH. Rabbits were selected for monoclonal antibody development based on reactivity in ELISA and Western blot assays. Four clones were selected for inclusion in the immunoenrichment kit (PTMScan^®^ O-GlcNAc [GlcNAc-S/T] Motif Kit #95220, Cell Signaling Technology, Inc.) to provide the broadest coverage of O-GlcNAc sites.

### Sample Generation

Mouse mESCs were routinely passaged by standard methods using 0.5% trypsin and ESC media (KO-DMEM, 10% FBS, 2 mM glutamine, 1X non-essential amino acids, 0.1 mM b-mercaptoethanol and recombinant leukemia inhibitory factor). Synaptosomal preps were prepared as previously described (30). Briefly, ten mouse brains were homogenized in homogenization buffer (5 mM HEPES (pH 7.4), 1mM MgCl2, 0.5mM CaCl2) supplemented with phosphatase inhibitors (PhosStop, Roche), protease inhibitors (cOmplete mini, Roche) and 10 µM PUGNAc using a Teflon homogenizer. After ten minutes of centrifugation at 1,400 x g, the supernatant was centrifuged at 13,800 x g for ten minutes. The pellet was resuspended in 0.32 M Tris-buffered sucrose and ultra-centrifuged into 1.2, 1.0, 0.85 M sucrose gradient at 82,500 x g, two hours. Synaptosome fractions between the 1 and 1.2 M sucrose layer were collected, pelletized and stored at −80°C for downstream processing. All steps were performed with ice cold buffers and centrifugation steps were performed at 4°C.

### Sample Preparation

Samples were prepared by two different methods. For urea-based digestion, cell pellets were resuspended in a 8M urea lysis buffer solution containing 75 mM NaCl, 50mM Tris HCl (pH 8.0), 1mM EDTA, Aprotinin (2 μg/μL) (Sigma-Aldrich), Leupeptin (10 μg/μL) (Roche), PMSF (1mM) (Sigma-Aldrich), Phosphatase Inhibitor Cocktail 2 (1:100 dilution of commercial stock, Sigma-Aldrich), Phosphatase Inhibitor Cocktail 3 (1:100 dilution of commercial stock, Sigma-Aldrich), and NaF (10mM). Protein concentrations were determined with a bicinchoninic acid (BCA) protein assay (Pierce). Lysis buffer solution was used to dilute protein concentrations to 8 mg/ml. Proteins were then reduced with 5 mM dithiothreitol (DTT) while shaking for one hour at room temperature. Proteins were subsequently alkylated with 10 mM of iodoacetamide for 45 minutes at room temperature shaking in the dark. Before digestion, proteins were diluted with 50 mM Tris HCl (pH 8.0) until the urea concentration was lowered to 2 M. Proteins were then digested for two hours with LysC (Wako Laboratories) for 2 hours at room temperature. Following this, proteins were digested with sequencing-grade trypsin (Promega) at room temperature while shaking overnight. Enzymatic digestion for both LysC and trypsin was performed at a 1:50 enzyme-substrate ratio. After digestion, peptides were acidified through the addition of formic acid (FA) to 1% and insoluble material was pelleted by centrifugation at 4,000 *x g* for 10 minutes. Peptides were desalted with 500 mg reversed-phase tC18 Sep-Pak cartridge (Waters). Cartridges were conditioned with 5ml of ACN, 5ml of 50% ACN 0.1% FA and equilibrated four times with 5mL of 0.1% trifluoroacetic acid (TFA). Peptides were loaded onto Sep-Pak cartridges, washed three times with 0.1% TFA, once with 1% FA, and then eluted with 50% ACN plus 0.1% FA. Peptide solutions were frozen, concentrated by vacuum centrifugation, and then resuspended in 30% ACN 0.1% FA. Peptide concentration was assessed via a BCA assay. Peptides were then aliquoted, frozen, dried to near completion, and stored at −80°C.

For SDS-based lysis, cell pellets or synaptosome preps were resuspended in 1 or 3% SDS, respectively, in a lysis buffer solution containing 50 mM TEAB, 1 mM EDTA, Aprotinin (2 μg/μL), Leupeptin (10 μg/μL), PMSF (1mM), Phosphatase Inhibitor Cocktail 2 (Sigma, 1:100 dilution of commercial stock), Phosphatase Inhibitor Cocktail 3 (Sigma, 1:100 dilution of commercial stock), and NaF (10 mM). Samples were sonicated using a Misonix Ultrasonic Liquid Processor (Fisher Scientific) probe sonicator pulsed twice for four seconds at an amplitude of 30 before a ten minute, 4°C centrifugation at 20,000 *x g*. Protein concentrations were assessed by BCA, reduced with 5 mM DTT for one hour, and subsequently alkylated with 10 mM iodoacetamide for 45 minutes in the dark while shaking at room temperature for both reactions. Proteins were precipitated using suspension trapping (31,32) using S-Trap midi columns (Protifi) per manufacturer’s recommendations. After digestion with trypsin, peptides were eluted, dried by vacuum centrifugation, and resuspended in 30% ACN 0.1% FA aqueous solutions. After quantification by BCA, peptides were aliquoted in their desired amounts, frozen, dried by vacuum centrifugation, and stored at −80°C.

### Enrichment of O-GlcNAc-modified peptides by Immunoprecipitation

Peptides were resuspended in chilled immunoaffinity purification (IAP) buffer at 1 mg/mL. Anti-O-GlcNAc protein-A coated agarose bead aliquots (PTMScan^®^ O-GlcNAc [GlcNAc-S/T] Motif Kit #95220Cell Signaling Technology, Inc.) were washed twice and resuspended in 160 uL of chilled IAP buffer. A range of reagent amounts were used (0.125x to 1x the recommended amount) for the antibody titration experiment. For all subsequent experiments, 0.25x (40 uL) of the resuspended antibody aliquot was used. Anti-O-GlcNAc Ab-beads were incubated with peptide samples for two hours at 4°C with end-over-end tumbling. After incubation, beads were pelleted by centrifugation for one minute. Flow throughs were transferred to separate tubes and frozen. Beads were first washed once with chilled IAP buffer and then twice with chilled PBS. Samples were eluted through resuspension with 150 uL of a 0.15% TFA aqueous solution with light manual agitation for five minutes. Samples were centrifuged for one minute and eluents were transferred to equilibrated Stage tips. The elution procedure was repeated twice before continuing with Stage tip sample clean up.

For all glycopeptide enrichment sample clean up, Stage tips (33) were prepared by stacking two punches of Empore 2215-C18 Octadecyl 47 mm disks (Bioanalytical Technologies) at the bottom of a 200 μL pipette tip. Stage tips were washed with 100 μL of methanol, 100 μL of 50% ACN 0.1% FA, and were equilibrated with 100 μL of 0.1% FA twice. Samples were loaded onto Stage tips after equilibration, washed twice with 100 μL of 0.1% FA and eluted with 50 μL of 50% ACN with 0.1% FA. Samples were then transferred to HPLC tubes, frozen, and dried by vacuum centrifugation. Samples were then resuspended in 9 μL of solvent A (3% ACN 0.1% FA) before LC-MS/MS analysis.

### Enrichment of glycopeptides with WGA

WGA glycopeptide enrichment was performed on 1 mg and 125 μg of urea digested mESC peptide aliquots in duplicate. Peptide samples were first resuspended in chilled IAP at 1 mg/ml. Next, 600 μL of WGA Agarose slurry (Vector Labs) stock solution was washed twice and resuspended in 1.2 mL of chilled IAP buffer. 400 μL of the washed WGA slurry was transferred to their appropriate labeled 1.5 mL tubes, and their respective peptide samples were then transferred. Samples were incubated at 4°C with end-over-end tumbling. After incubation, samples were pelleted by centrifugation for one minute. Flow throughs were transferred to separate tubes and preserved. Beads were first washed once with chilled IAP buffer and then twice with chilled PBS. Peptides were then eluted by resuspending samples in 300 μL of 0.15% TFA. The resuspended samples were then immediately transferred to the top of equilibrated Stage-tips and allowed to trans-elute onto C18 disks for 5 minutes. Stage tips were washed once with 75 μL of 0.1% FA and eluted with 50 ul of 50% ACN with 0.1% FA. Samples were transferred to HPLC vials, frozen, and dried by vacuum centrifugation. Samples were resuspended in 9 μL of solvent A (3% ACN 0.1% FA).

### Basic Reversed-Phase (bRP) Chromatography

Four mg of S-trap digested synaptosome peptides were fractionated by basic reversed-phase (bRP) chromatography. bRP solvent A was made in an aqueous solution of 5mM of Ammonium Formate and 2% ACN (pH 10). bRP Solvent B was made in a solution of 5 mM Ammonium Formate and 90% ACN (pH 10). A 4.6 mm inner diameter 250 mm long Agilent Zorbax 300 Extend-C18 (Agilent) was used as the bRP column. Chromatography occurred over a 96-minute gradient with fractions collected in a Whatman 2ml 96 well plate (GE Healthcare). A 1260 Infinity II high-performance liquid chromatography (HPLC) instrument was used (Agilent). The chromatography gradient proceeded with a solvent B composition scheme of: 0% to 5 minutes, 5% at 7.66 minutes, 28.5% at 64.98 minutes, 34% at 70.48 minutes, 60% at 83.48 minutes, and 60% solvent B continuing for the remainder of the gradient. Peptides were concatenated into a 12-fraction scheme as previously described (34). Peptides were then frozen and dried by vacuum centrifugation in preparation for O-GlcNAc enrichment.

### Mass Spectrometric Data Acquisition

Peptides were separated online using an Easy-nLC 1200 HPLC (Thermo Fisher Scientific). Samples were loaded onto a capillary column with an integrated emitter tip and heated to 52°C. The capillary column was packed in-house with ReproSil-Pur C18-AQ 1.9 μm beads (Dr. Maisch GmbH) to an approximate length of 27 cm. Once loaded onto the capillary column, peptides were separated through acidic reversed-phase chromatography with solvent A composed of 3% ACN and 0.1% FA and solvent B composed of 90% ACN and 0.1% FA. Chromatography proceeded for 110 minutes with the following gradient of solvent B composition: 2% solvent B at the start, 6% at 1 minute, 30% at 85 minutes, 60% at 94 minutes, 90% at 95 minutes, 90% at 100 minutes, 50% at 101 minutes, and 50% solvent B continuing for the remainder of the gradient. Chromatography had a flow rate of 200 nL/min until 101 minutes, at which the flow rate increased to 500 nL/min for the remainder of the gradient.

Data was acquired on a Thermo Scientific Orbitrap Fusion Lumos mass spectrometer. 4/9 μL of sample was injected for each LC-MS/MS run, with the exception of the HCD collision energy (CE) ramp experiment where 3 μL was injected per run. All MS1 scans were performed in the Orbitrap at 60K resolution with a scan range of 350-1800 m/z with a maximum injection time of 50 milliseconds. Unless otherwise specified, all samples were assessed using the HCD-pd-ETD method for MS2 data acquisition. HCD scans containing either a 204.0867 m/z or 138.0545 m/z peak within 15 ppm of the top 20 most intense peaks would trigger a separate ETD scan on the same precursor. All ETD in this study was performed with 25% HCD supplemental activation (EThcD). MS2 scans were analyzed in the Orbitrap for both HCD and EThcD at 15,000 resolution and 1.7 m/z isolation window, with 105 millisecond injection time and normalized AGC of 200% for HCD and a 120 ms injection time with a normalized AGC of 800% for EThcD. HCD was set at 30 CE for antibody and peptide titration experiments. HCD CE was set to 40 for the 12 fraction synaptosome O-GlcNAc enriched samples. A two second cycle duty time was set for MS/MS events, with monoisotopic precursor determination set to “Peptide”, a filter intensity threshold set to 1.0e4, a charge state filter between 2-8, and a 30 second dynamic exclusion time. Precursor priorities were set to highest charge state and lowest m/z.

Data for EThcD alone was acquired by reinjection of the 12 fraction O-GlcNAc enriched synaptosome samples. MS1 parameters remained the same as previously described. EThcD was conducted with 25% supplemental HCD activation, analyzed in the Orbitrap at a resolution of 15,000 with an isolation window of 1.7, a normalized AGC target of 60%, and a 50-millisecond injection time. The duty cycles was set to 20, with monoisotopic precursor determination set to “Peptide”, a filter intensity threshold set to 1.0e4, a charge state filter between 2-6, and a 45 second dynamic exclusion time.

### Data Analysis

Raw files were processed and searched using Byonic (Protein Metrics Inc.). Cleavage specificity was set to RK with C-terminal cleavage and “Fully Specific” specificity. Only two missed cleavages were allowed. Precursor and product ion mass tolerances were set to 10 and 20 ppm, respectively. Cysteine carbamidomethylation was set as a fixed modification, where methionine oxidation was set to variable and “common2”. Protein N-term acetylation was set to “rare1”. HexNAc (N-acetylhexosamine) at S and T was set to “common3” and phospho at S, T, and Y was set to “common2”. A total of four common and one rare modification was allowed. Peptide identifications were extracted using Byos (Protein Metrics) for each run individually and combined into a single report for further processing. PSMs were excluded if they did not include one HexNAc mass addition, or fell below a two-dimensional posterior error probability (PEP2D) cutoff of 0.05. “Localized HexNAc PSMs” were putatively identified by filtering out all HexNAc PSMs with a Delta Mod Score greater than or equal to ten (35). Distinct glycopeptides were determined differently for HCD than for EthcD in the synaptosome analysis. For HCD-based identifications site assignments forced by Byonic’s output were removed from the glycopeptide sequence and concatenated with the number of HexNAcs determined by the precursor mass. Sequence_#HexNAcs were then collapsed, retaining the PSM with the highest Delta Mod Score, and are referred to as “Distinct glycopeptides”. For EThcD - based identifications, the glycosylation site(s) within the peptide was retained, and PSMs were collapsed retaining the PSM with the highest Delta Mod Score. Distinct glycopeptides for both fragmentation methods were determined in the same way for the reproducibility analysis, where site assignments forced by Byonic’s output were removed from the glycopeptide sequence, concatenated with the number of HexNAcs determined by the precursor mass, and collapsed down to unique entries.

Motif analysis was performed by centering each distinct glycosylation site with a Delta Mod Score greater than or equal to ten, with five flanking amino acids. Sequence logos were created by Weblogo (https://weblogo.berkeley.edu/logo.cgi) using default parameters, and by PhosphositePlus (https://www.phosphosite.org/staticMotifAnalysis#) using the Automated Selection as background. Identification overlap analysis was performed using distinct glycopeptides or distinct Uniprot accession numbers/proteins from each run.

### Quantification of Oxonium Ion-containing Spectra

MS2 peak lists were generated for 1 mg mESC peptide WGA & Ab-based enrichments. Spectral .raw files were first processed in Spectrum Mill version BI.07.04.210 (Broad Institute of MIT & Harvard) using the Extractor feature to obtain .mzXML files. Spectral filtering set between 600-6000 Da mass range for precursors and a sequence tag length of greater than 0. The instrument setting was set to ESI Q-Exactive HCD and carbamidomethylation on cysteines was set as a fixed modification. Spectra from the same precursor or within a retention time window of +/− 60 seconds were merged.

Extracted files were uploaded to the MS-Filter feature in Protein Prospector v 6.2.2 (UCSF Mass Spectrometry Facility). Precursor mass range was set to 600-6000 Da, charge filter was set to 2-8, masses were set to be monoisotopic, parental tolerances were set to 200 ppm, fragment tolerances were set to 20 ppm, an ESI-Q-high-res instrument setting was selected, and Max MSMS Pks was selected.

Peak lists were exported from Protein Prospector as .txt files. Files were uploaded to a Python script using the pandas library. The Python script was developed to convert peak lists into dataframes where scan numbers were row identifiers, predetermined peaks were column identifiers (366.140, 204.087, 186.076, 168.066, 144.065, 138.055, 126.055, & 274.092 within a tolerance of 20 ppm), and dataframe values were peak intensities. A value.count() function was applied to these data frames to obtain the number of spectra (MS2 events) with 204, 366 & 274 peaks.

### Distribution of Ratios for Oxonium Ion Fragments

Applying the Python script described earlier, dataframes for glyco-diagnostic ion containing MS2 spectra were created for CD43, MUC16, MUC16 treated with sialidase, MUC16 treated with sialidase and PNGaseF, and a 1 mg mESC peptide Ab-based enrichment. A GlcNAc/GalNAc ratio column was created by taking the sum of the 138 and 168 peaks divided by the sum of the 126 and 144 peaks for MS2 spectra with those four peaks present. The distribution of this ratio was plotted via a kernel estimate density using distplot() function in the seaborn library (**Figure 3A**). The same strategy was also applied for the 138 / 144 peak ratios (**Figure 3B**). Outlier values, as defined by values three z-scores away from the mean, were removed for the plotted ratio columns. Computational removal of HexHexNAc and sialic acid was performed by subsetting data frames to contain row indices without values in either the 366.140 or 274.092 columns.

### Preparation of CD43 and MUC16 for Oxonium Ion Comparison

Samples were prepared as previously described (36,37). Briefly, recombinant glycoproteins (R&D Systems 1658-PD and 3345-PS) were digested overnight with mucinase SmEnhancin (38) at 37°C in a total reaction volume of 12 μL in 50 mM ammonium bicarbonate. In sialidase treated samples, 1 μL of dilute (1:40) sialoEXO (Genovis) was also added to the mucinase digestion. The volume was then increased to 19 μL with 50 mM ammonium bicarbonate. For PNGaseF treated samples, 1 μL of enzyme (Promega) was added to 99 μL of 50 mM ammonium bicarbonate, and 1 μL of this reaction was added to each mucinase reaction vial. De-N-glycosylation reactions were incubated for 8-12 h at 37°C. For reactions without PNGaseF, 1 μL of ammonium bicarbonate was added. Reduction and alkylation were performed according to ProteaseMax (Promega) protocols. A total of 1 μL of 0.5 M DTT was added and the samples were incubated at 56°C for 20 min, followed by the addition of 2.7 μL of 0.55 M iodoacetamide at room temperature for 15 min in the dark. Digestion was completed by adding sequencing-grade trypsin (Promega) at a 1:20 E:S ratio overnight at 37°C and quenched by adding 0.3 μL of glacial acetic acid. C18 clean-up was performed using 1 mL strataX columns (Phenomenex). Each column was incubated with 1 mL of acetonitrile once, followed by one 1 mL rinse of buffer A (0.1% formic acid in water). The samples were diluted to 1 mL in buffer A and loaded through the column, then rinsed with buffer A. Finally, the samples were eluted with three rinses of 100 μL of buffer B (0.5% formic acid, 80% acetonitrile) and dried by vacuum centrifugation. The samples were reconstituted in 10 μL of buffer A for MS analysis. Samples were analyzed on an Orbitrap Fusion Tribrid (Thermo) using an HCD-pd-ETD method, as described above, with no supplemental activation for ETD.

## Results

### Anti-O-GlcNAc Antibodies Provide a Simple, Sensitive Approach to Enrich Native O-GlcNAc-modified Peptides

Four new anti-O-GlcNAc rabbit monoclonal antibodies (Abs) were recently developed by Cell Signaling Technology^®^ and are provided as a proprietary mixture coupled to Protein A-coated agarose beads (PTMScan O-GlcNAc [GlcNAc-S/T] Motif kit #9522). To initially characterize the performance of this Ab-based enrichment reagent, we generated tryptic digests of whole cell lysates (WCL) from mouse embryonic stem cells (mESCs). mESCs were chosen due to their relatively high levels of O-GlcNAc, and because OGT is essential for embryonic development (1,2,39). Data were acquired on a Fusion Lumos Tribrid mass spectrometer using an HCD product-triggering EThcD (HCD-pd-EThcD) acquisition method. HCD was performed on all multiply charged precursor ions. Upon detection of the glyco-specific oxonium fragment ion masses of 204.087 or 138.055 (+/− 15 ppm) in the HCD scan, the precursor was re-accumulated for a separate ETD scan with supplemental HCD (25%) activation (EThcD) (9,15). As HexNAc oxonium ion detection in the HCD scan is crucial to trigger the subsequent EThcD scan necessary for confident O-HexNAc (O-linked N-acetylhexosamine) site assignment, we first tested various HCD collision energies (CEs) using WCL digests of mESC enriched using the anti-O-GlcNAc Ab mixture. We found that higher HCD CEs caused a slight decrease in the number of HexNAc-containing HCD peptide spectral matches (PSMs) but increased the number of HexNAc-containing EThcD-derived PSMs (**Figure 1A**). Higher HCD CEs also resulted in higher spectral quality for both HCD and the product-triggered EThcD (pd-EThcD), measured by the improved confidence (PEP2D score) in glycopeptide identifications (**Figure 1B**). We therefore chose these data acquisition conditions for analysis of O-GlcNAcylated peptides.

**Figure 1.**
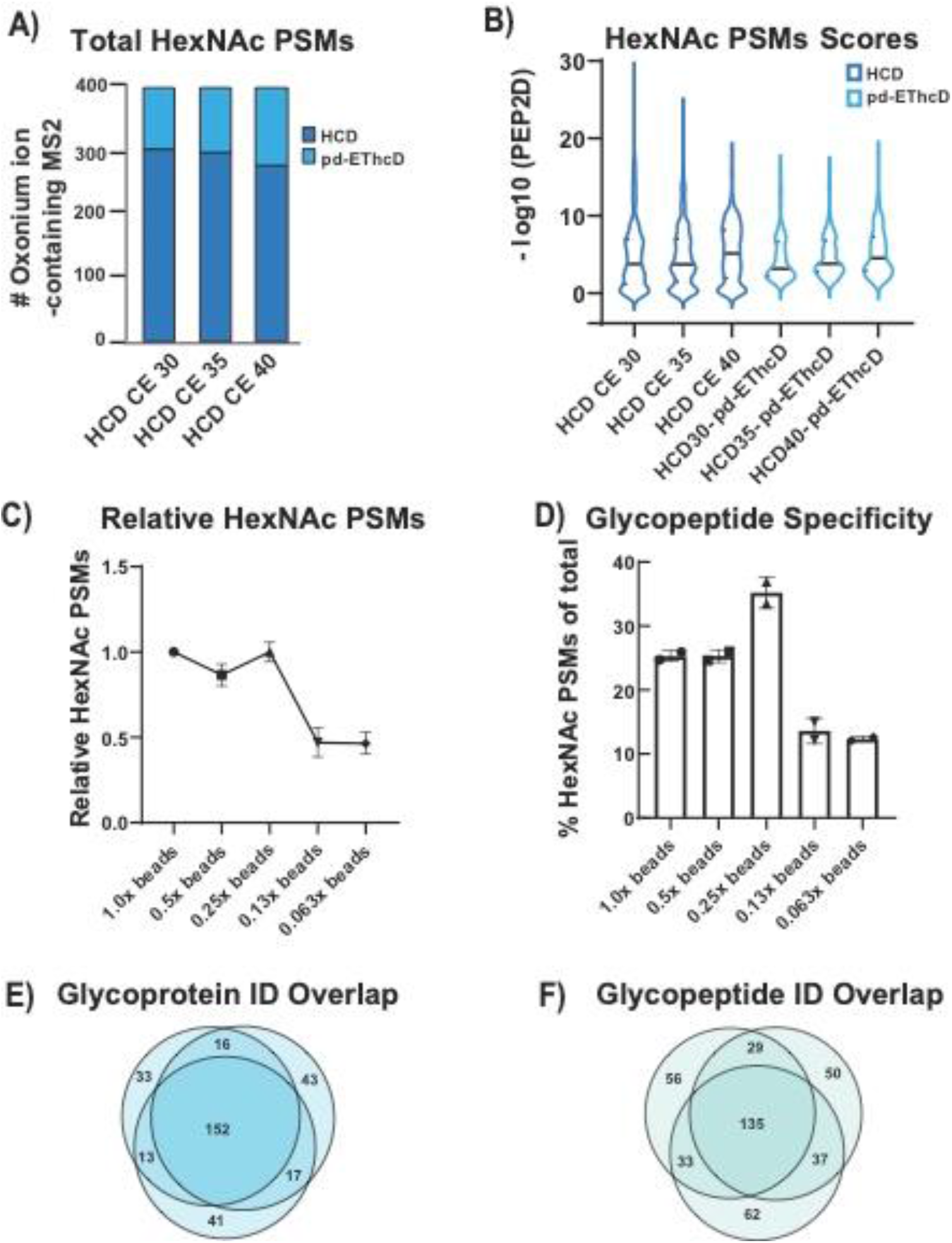
Optimization of Data Acquisition and Antibody Enrichment Conditions **A)** Total HexNAc-containing PSMs across different HCD collision energies (CEs) (300 ug of mESC WCL). Both HCD and product triggered EthcD (pd-EthcD) PSMs are shown. Only HCD CEs were altered. **B)** Distribution of PEP2D scores for HexNAc-containing PSMs separated by dissociation method. Only HCD CEs were altered. **C)** Relative HexNAc-containing PSMs titrating down the amount of antibody from the recommended levels. Values are relative to the 1x experiment. **D)** O-HexNAc glycopeptide specificity for decreasing amounts of antibody. Specificity was calculated by dividing the number of HexNAc-containing PSMs by the total number of PSMs. **E)** Overlap in distinct glycoprotein identifications from triplicate 1 mg mESC WCL enrichments. **F)** Overlap in distinct glycopeptide identifications from three separate enrichments from 1 mg aliquots of digested mESC WCL digests.

We next optimized the amount of Ab-conjugated beads needed to observe the highest number of HexNAc-containing PSMs and the lowest number of unmodified PSMs using one milligram of mESC WCL digest. We found that use of one-fourth the amount of Ab recommended for O-GlcNAc-peptide yielded equivalent numbers of HexNAc-containing PSMs as the full amount while providing the best enrichment specificity (**Figure 1C-D**), consistent with previous anti-PTM Ab work (40,41). Reproducibility was evaluated across three parallel enrichments of the same mESC sample. We found the overlap of glycoprotein identifications to be 48% across all three replicates or an average of 60.63% across any two (**Figure 1E**). At the distinct glycopeptide level, the overlap across all three enrichments was 34%, or an average of 49% for across any two (**Figure 1F**). This level of reproducibility across replicated is similar to that obtained using anti-phosphotyrosine antibodies (42).

To evaluate the sensitivity of the anti-O-GlcNAc Abs, we enriched from decreasing amounts of input peptides, keeping the amount of Ab-beads constant. We observed a nearly linear decrease in HexNAc-containing PSMs as input decreased, where the greatest drop occurred between 125 and 250 ug peptides (**Figure 2A**). The site, peptide and protein identification results of the 1 milligram enrichment are attached in **Supplemental Table 1**. We compared these results to an alternative, commonly used native O-GlcNAc-modified peptide enrichment reagent WGA (23,24,26,43), at our high and low input loads. In batch mode enrichments, the anti-O-GlcNAc Abs greatly improved the number of HexNAc-containing PSMs across all input levels compared to WGA (**Figure 2A**). Separating the HexNAc-containing PSMs by unimolecular dissociation mode showed similar trends, illustrating that although HCD provides little site assignment information it is responsible for roughly two-thirds of PSM identifications (**Figure 2B**).

**Figure 2.**
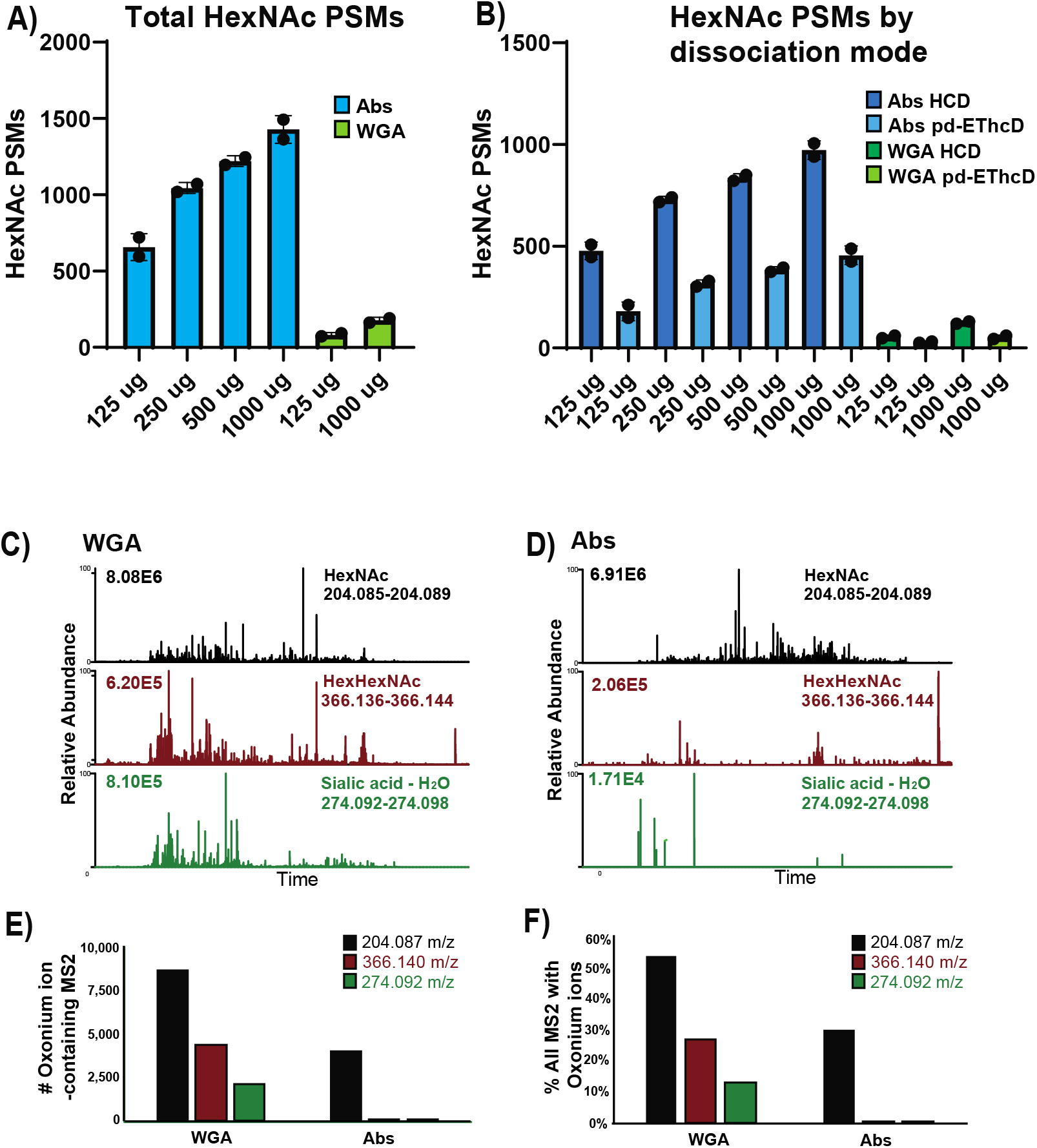
Characterization of sensitivity and specificity of anti-O-GlcNAc antibodies and their comparison with WGA **A)** Total number of HexNAc-containing PSMs obtained by enrichment with Ab (blue bars) or WGA (green bars) across a range of input levels of mESC WCL digest; the optimal amount of Ab determined in Figure 1 was used. **B)** Numbers of HexNAc-containing PSMs obtained at varying input levels and separated by dissociation method. “pd-EThcD” indicates the EThcD scan was triggered by detection of HexNAc oxonium ions in the preceding HCD scan. **C)** Extracted Ion Chromatograms (XICs) from the WGA-based glycopeptide enrichment experiments of oxonium ions (+/− 20 ppm of calculated masses) diagnostic for respective carbohydrates; 204.087 m/z represents HexNAc (black), 366.140 m/z for HexHexNAc (burgundy), and 274.092 m/z for sialic acid minus water (green). Base peak intensities are shown on the upper left. **D)** Same as in C, above, but for the anti-O-GlcNAc Ab-based glycopeptide enrichments. **E)** Number of MS2 spectra containing one of the aforementioned oxonium ions from the respective enrichment strategies from 1 mg of peptide inputs. **F)** Percentage of all MS2 scans with aforementioned oxonium ions from the respective enrichment strategies from 1 mg of peptide inputs.

### Anti-O-GlcNAc Antibodies are highly specific for O-GlcNAc-modified Peptides

To evaluate the specificity of the anti-O-GlcNAc Abs for O-GlcNAc-modified peptides we compared the results obtained with the Abs to the use of wheat germ agglutinin (WGA), a lectin with affinity for GlcNAc and a diverse array of glycosylation states (28). The extracted ion chromatograms (XICs) of the WGA-based enrichment runs showed a high prevalence of diagnostic ions for HexNAc (204.087 m/z), HexHexNAc (366.140 m/z), and sialic acid minus water (SA, 274.092 m/z) (**Figure 2C**). The diagnostic ions of m/z 366 and 274 are derived from extended glycan structures (e.g., biantennary and more highly branched and modified oligosaccharides) on peptides and likely not from an OGT substrate (44). Whereas, the observation of the 204 m/z peak in the absence of 366 or 274 is suggestive of an O-linked HexNAc and not extended carbohydrate structures. In contrast to the WGA results, XICs from the Ab-based enrichment showed a high number of m/z 204 HexNAc peaks with few HexHexNAc or SA diagnostic ions (**Figure 2D**). Additional evidence of specificity of the Abs for O-GlcNAc was obtained by quantifying the number of MS2 scans that exhibited the respective diagnostic oxonium ions (**Figure 2E**). While the WGA-based enrichments had twice as many HexNAc-containing spectra than the Ab-based enrichments, they also contained 91- and 264-fold more HexHexNAc- and SA-containing spectra, respectively, compared to the Ab-based enrichments. The percentage of total MS2 spectra with the respective diagnostic ions showed similar trends (**Figure 2F**). These data suggest that the anti-O-GlcNAc Abs have a high degree of specificity for non-extended O-HexNAc-modified peptides and do not enrich HexNAc-containing complex glycans to a significant extent.

While the anti-O-GlcNAc monoclonal Abs were developed against synthetic O-GlcNAc-modified peptides with degenerative amino acid sequences, we questioned whether the HexNAc signal we observed was coming from non-specific enrichment of O-linked N-acetylgalactosamine (O-GalNAc)-modified peptides. Singly O-GalNAc-modified serines and threonines (a.k.a. Tn antigen) is the base of the Core 1 O-glycosylation pathway in the Golgi (Reviewed in (45)). GlcNAc and GalNAc are isobaric, displaying the same oxonium ion and fragments thereof, making these two species challenging to differentiate by mass alone. In work analyzing synthetic glycopeptides by HCD-MS/MS, other groups have suggested that O-GlcNAc and O-GalNAc undergo different dissociation pathways and the ratio of the resultant oxonium ions can be used to discriminate between these epimers (21,46). We developed a script to calculate the ratio of select oxonium ions for each individual HCD MS2 spectrum, focusing on the GlcNAc/GalNAc ratio suggested by Halim *et al*., [(138 + 168)/(126 + 144)] (46). To test the ability of our script to differentiate between GlcNAc and GalNAc oxonium ions, we turned to proteins with well-characterized glycosylation states for digestion and analysis by HCD-MS/MS. For a model that is predominantly O-GalNAcylated, we used the mucin SPN (a.k.a. CD43 or Leukosialin) which has 60 documented O-GalNAc sites, and a single annotated N-linked glycosylation site (47). For a protein containing large numbers of both O-GalNAc and N-linked glycans, we used the SEA domain of human MUC16. Plotting the GlcNAc/GalNAc ratios for these two proteins, we found that SPN, the highly O-GalNAc-modified protein, showed a tight distribution of ratios with a mean of 0.8. MUC16 had a mean of 3.3 and a much broader distribution, likely due to its diverse range of glycosylation types (**Figure 3A, SPN, orange line; MUC16, green line**). Consistent with Halim *et al.*, the large GlcNAc/GalNAc ratios of MUC16 were likely due to N-linked glycans containing GlcNAc residues. To further test this hypothesis, we removed specific sugars from MUC16 in multiple ways. First, we treated MUC16 with sialidase, which removes terminal SA, and replotted the GlcNAc/GalNAc ratio. Sialidase treatment increased the number of lower ratios but kept the same wide distribution as untreated MUC16. We next removed the N-linked glycans from MUC16 with PNGase F, which cleaves glycans from Asn but not Ser or Thr. The GlcNAc/GalNAc ratios from PNGase F-treated MUC16 greatly increased the number of low ratios and decreased the number of high ratios, creating a distribution resembling SPN’s glycopeptides. Finally, we computationally removed MS2 spectra from MUC16 that contained either 366.140 or 274.092 m/z to simulate a perfect enzymatic reaction that should leave only O-GalNAc-modified peptides. The GlcNAc/GalNAc distribution of this “computational glycosylation removal” showed a similar pattern to that of the PNGase F-treated MUC16 and the heavily O-GalNAc-modified SPN. These results show our ratio distribution analysis can discriminate between O-HexNAc and hybrid, complex glycans.

**Figure 3.**
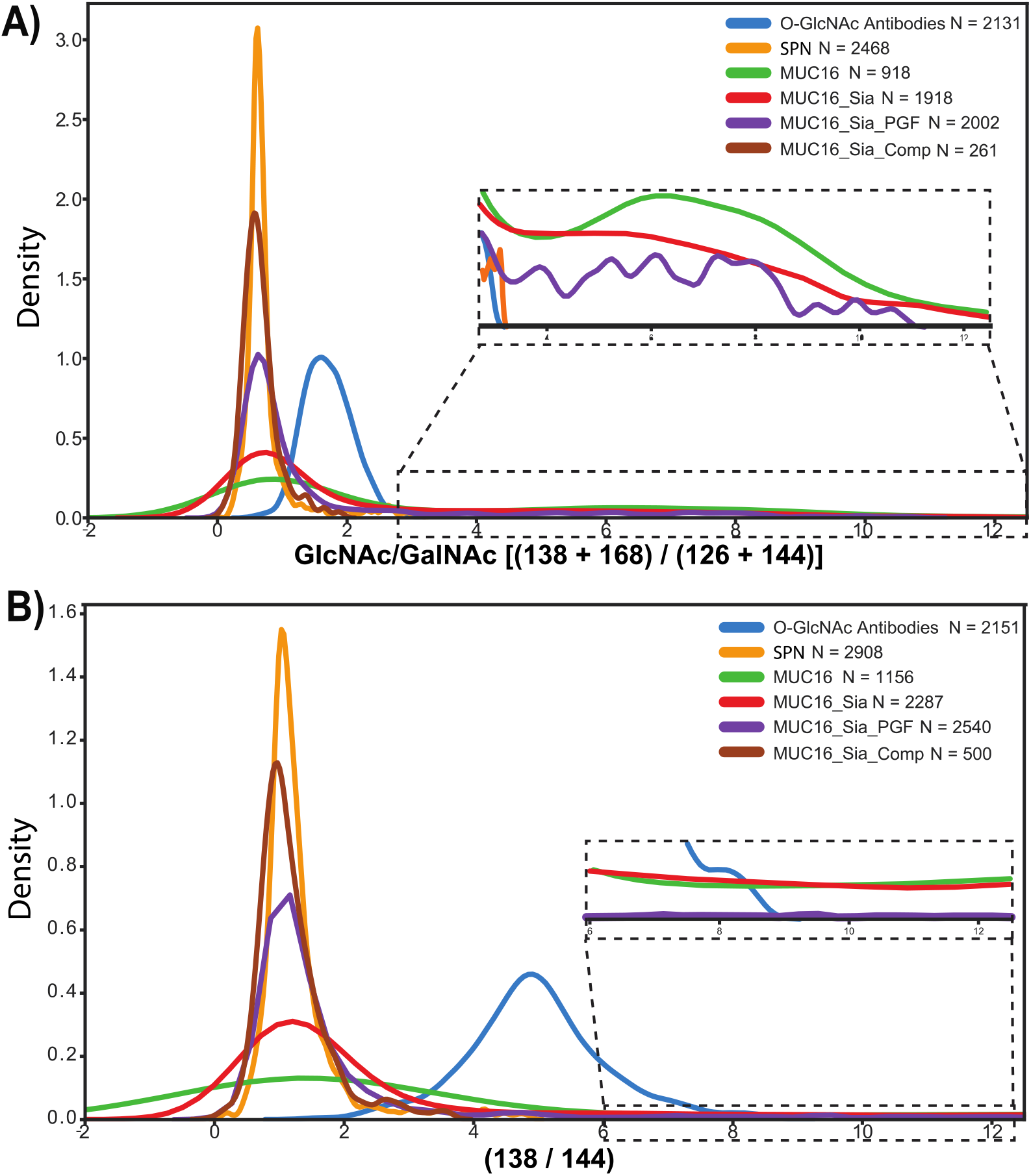
Discrimination between GlcNAc and GalNAc through Oxonium Ion Ratio Distributions **A)** Kernel density plots for the distribution of GlcNAc/GalNAc ratios [(138 + 168) (126 + 144)], as determined by Halim *et al. (46)* for control proteins and conditions. Inset is a zoom of the higher ratios for all values highlighted. N = the number of MS2 spectra used to derive ratios. Sia = sialidase treatment; PGF = PNGase F treatment; Comp = computational removal of MS2 spectra containing diagnostic ions for 366 and 274. **B)** Kernel density plots for the distribution of 138 144 peaks across the same samples in Figure 3A. Inset is a zoom of the higher ratios. Abbreviations are the same from Figure 3A.

To test whether our GlcNAc/GalNAc distribution analysis could differentiate between O-GalNAc and our putative O-GlcNAc-modified peptides, we plotted the GlcNAc/GalNAc ratios for MS2 spectra generated from the anti-O-GlcNAc Ab-based enrichments. We found that the GlcNAc/GalNAc ratios from these samples were distinctly higher (mean = 1.7) than those from SPN and had a tighter distribution than those derived from MUC16 (**Figure 3A**). Consistent with Halim *et al*., our putative O-GlcNAc-containing spectra had intermediate ratios compared to O-GalNAc and more complex glycans derived from MUC16, suggesting we are indeed enriching O-GlcNAc-modified peptides. To better discriminate between O-GlcNAc and O-GalNAc, we examined alternate oxonium ion ratio distributions. Our data showed that using a simpler ratio (138 m/z / 144 m/z) provided stronger separation between O-GalNAc-only spectra and the data from our Ab-based enrichments (**Figure 3B**). These results demonstrate that the ratio of the 138 / 144 oxonium ions can differentiate between O-GlcNAc and O-GalNAc. Together, these data suggest that the anti-O-GlcNAc Abs show strong specificity towards O-GlcNAc, and not O-GalNAc or GlcNAc residing in more complex glycan-modified peptides.

### Anti-O-GlcNAc antibodies allow for deep O-GlcNAc proteome profiling from tissues

An advantage of native O-GlcNAc-modified peptide enrichment over alternative strategies such as metabolic labeling is the ability to enrich glycopeptides from tissues. To test the capacity of the anti-O-GlcNAc Abs to enrich from tissues, we turned to synaptosome preparations (30), a classic model used for O-GlcNAc-modified peptide enrichment (12,24,43). Murine synaptosomes were solubilized in 3% SDS, subjected to suspension trapping (31,32), and digested with trypsin. Four milligrams of the synaptosome digest were fractionated with basic reversed-phase (bRP) fractionation into 96 fractions then concatenated back to 12 fractions (34). Each final fraction containing approximately 330 μg of peptide was enriched for O-GlcNAc-modified peptides using the anti-O-GlcNAc Abs, and the data was acquired using either the HCD-pd-EThcD method or EThcD alone without a preceding HCD scan.

For both data acquisition methods, the number of HexNAc-containing PSMs was fairly evenly spaced across the twelve fractions with the exception of fraction one (**Figure 4A**). Using the HCD-pd-EThcD acquisition method we identified 6,872 high confidence HexNAc-containing PSMs, 4,427 of which were identified by HCD and 2,445 by pd-EThcD (**Figure 4B**). After collapsing redundant HexNAc PSMs, 1,328 distinct glycopeptides remained by HCD and 1,289 distinct glycopeptides by pd-EThcD. Of the distinct glycopeptide forms identified by pd-EThcD, 714 had a Delta Mod Score of 10 or greater, the suggested lower limit of confidence for site assignment by Byonic (35). Using this metric, we identified 1,040 O-HexNAc sites through pd-EThcD (**Supplemental Table 2**). While HCD of O-linked glycopeptides generally does not retain site localization information due to the prominent cleavage of the O-glycosidic bond, we found 190 distinct glycopeptide forms with Delta Mods scores of 10 or greater, leading to 272 O-HexNAc sites. Site assignments with HCD were enabled largely by having fewer S/T residues within the peptides (**Figure 4C**). With EThcD acquisition alone we identified 1,532 high confidence HexNAc-containing PSMs containing 1,316 localized HexNAc PSMs (**Figure 4B**). After collapsing redundant glycoPSMs, EThcD alone identified 919 distinct glycopeptides with 791 O-HexNAc sites with a Delta Mod Score of ten or greater (**Supplemental Table 3**). Although HCD provides little O-HexNAc site information, its use led to twice as many distinct glycoprotein identifications compared to EThcD alone (**Figure 4D**). These data show that HCD provides complimentary information on glycopeptide and glycoprotein identifications to EThcD alone, without sacrificing the number of EThcD-based identifications.

**Figure 4.**
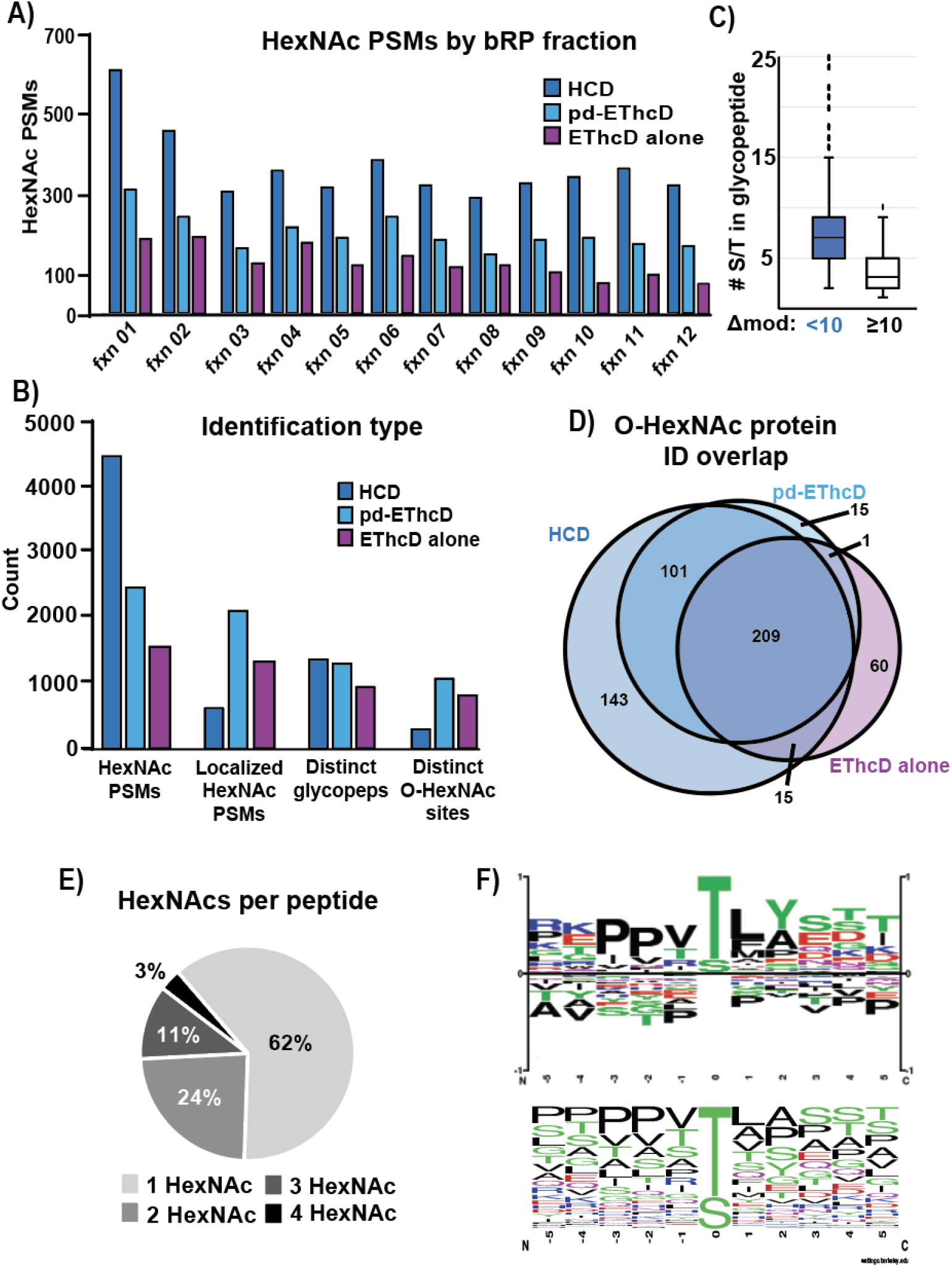
Benchmarking Anti-O-GlcNAc Antibodies with Post-Synaptic Density Preparations across Different Data Acquisition Methods **A)** HexNAc-containing PSMs across basic reversed-phase fractions. Blue bars indicate the number of HexNAc-containing PSMs identified by HCD or the subsequent pd-EThcD scan. Purple bars indicate the number of HexNAc-containing PSMs from EThcD alone. **B)** Breakdown of various identifications from the cumulative analysis analyzing O-GlcNAc peptides from mouse synaptosome fractions. “HexNAc PSMs” refer to PSMs that contain at least one HexNAc modification, regardless of localization. “Localized HexNAc PSMs” refers to all glycosylation sites within HexNAc-containing PSMs that had a Delta Mod Score of ten or greater. “Distinct Glycopeps” refers to non-redundant glycopeptides identified by the respective dissociation mode. However, HCD-based identifications were treated as if site localization could not be used (see Methods). “Distinct O-HexNAc sites” refers to the count of HexNAc site-assignments from all sites within non-redundant HexNAc-containing PSMs with Delta Mod Score of ten or greater. **C)** Number of S/T residues within HCD-based glycopeptide identifications with a Delta Mod Score of less than ten (blue box) or ten or greater (white box) to symbolize the trend HCD mostly identifies O-HexNAc sites in peptide with few amino acid possibilities for modification. **D)** Overlap of proteins identified as O-HexNAc modified by the respective data acquisition methods. Only a single HexNAc-containing PSM was required to be called an O-GlcNAc-modified protein. “pd-EThcD” indicates the EThcD scan triggered from the detection of HexNAc oxonium ions in the antecedent HCD scan. **E)** Distribution of the extent of HexNAc modifications of HexNAc-containing PSMs across all high-confidence HexNAc-containing PSMs. **F)** Sequence logo of distinct O-HexNAc sites according to two different analysis methods (see Methods). The top logo was derived from Phosphosite.org; the bottom from Weblogo (Berkeley).

To better understand the properties of O-HexNAc-modified peptides enriched using the anti-O-GlcNAc Abs, we analyzed the characteristics of the identified glycopeptides. We first looked at the extent of glycosylation per peptide. Of all HexNAc PSMs across both datasets, 62% had a single HexNAc per glycopeptide (**Figure 4E**). Interestingly, 38% of HexNAc PSMs were multiply O-HexNAcylated, roughly twice as many as seen previously (24). We next explored the frequency of amino acid occurrences in the sequence surrounding the O-HexNAc site. We used two commonly used motif analysis tools, Weblogo (48) and PhosphositePlus (49), as they both use different background corrections (see Methods). We found both sequence logo generators showed threonine to be more often O-HexNAc modified than serine (**Figure 4F**), a departure from previous studies (12,24,50,51). Consistent with previous work, we found a preference for proline residues at position −2 and −3, some hydrophobic character at position +1 and +2, and an enrichment for hydroxyl-containing residues C-terminal to the distinct O-HexNAc sites. Together these data suggest that the anti-O-GlcNAc Abs efficiently and specifically enrich for O-GlcNAc-modified peptides possessing the weak primary sequence preference of OGT.

## Discussion

Here we report the in-depth characterization and application of new anti-O-GlcNAc Abs to enrich for native O-GlcNAc-modified peptides from cells and tissues. We also describe the benefits and drawbacks to combining HCD with EThcD for the analysis of a complex mixture of O-GlcNAc peptides.

### Benefits and Drawbacks of HCD Product-Triggering EThcD

While the necessity of EThcD for O-linked glycopeptide analysis is well-established (13–15), the value of combining it with HCD for global O-GlcNAc proteome analysis proved manifold. The HCD-derived HexNAc-PSMs provided additional, complimentary evidence for peptide O-GlcNAcylation, though site localization was minimal. The speed of HCD also provided a substantial increase in O-GlcNAcylated peptide/protein identifications compared to EThcD alone. We saw little to no decrease in O-GlcNAc site information when using the HCD “scouting” scan to trigger EThcD. In fact, we found that increasing HCD collision energies led to higher scoring O-HexNAc-modified PSMs in both HCD and EThcD. This suggests that HCD “scouting” scan to trigger EThcD will be compatible with isobaric labeling for quantitation (iTRAQ, TMT) of O-GlcNAcylation (9,15,27) and is a subject of our future studies.

### Anti-O-GlcNAc Abs Provide a Simple, Efficient and Specific Enrichment Strategy for Analysis of Native O-GlcNAc Peptides from Cells and Tissues

We found the novel anti-O-GlcNAc Abs coupled to Protein A-coated agarose beads to be a strong enrichment reagent to analyze O-GlcNAc-modified peptides from cells and tissues. A major benefit of the anti-O-GlcNAc Abs is their specificity towards O-GlcNAc vs. O-GalNAc or extended, GlcNAc-containing glycans. We saw little evidence of HexHexNAc -or sialic acid-containing peptides immunoprecipitated by the Abs, suggesting they do not recognize GlcNAc within glycan chains. This specificity was reflected in the higher number of HexNAc-containing PSMs compared to WGA, where our search parameters did not allow for any glycan mass additions. We also found evidence for a strong specificity of the anti-O-GlcNAc Abs towards O-GlcNAc and not its epimer O-GalNAc. Leveraging prior studies (21,46), we confirmed that the distributions of the ratios of HexNAc oxonium ion fragments were distinct between a heavily O-GalNAc-modified protein and our putative O-GlcNAc-modified peptides. We then modified the previous GlcNAc/GalNAc ratio to utilize only two fragment ions, 138 / 144, which provided a clearer separation between the two glycopeptide forms. We suggest this ratio should be used in future studies to help differentiate between these isobaric molecules. Future work will be needed to understand how to use both these ratios, and potentially other oxonium ions and fragments thereof, to estimate the relative sugar composition of more complex glycans, especially when examining the higher ratios.

The simplicity of use and sensitivity of the anti-O-GlcNAc Abs are also major advances for global O-GlcNAc analysis. Compared to other landmark studies (12,24,43), we achieved similar numbers of distinct glycopeptides and O-GlcNAc sites, using nearly ten-fold less material, and a significantly smaller fraction of labor and instrument time. The sample preparation time is dramatically decreased compared to other O-GlcNAc enrichment protocols and can be done with little to no special equipment or chemistries. As this enrichment strategy does not require incubating live cells with chemically-modified sugars, it is compatible with enrichment from tissues, demonstrated by our analysis of synaptosomes. We expect that the enrichment of O-GlcNAc-modified peptides using these Abs can be easily coupled to serial PTM enrichment workflows routinely used in our laboratory (34).

In summary, we provide an in-depth characterization of novel anti-O-GlcNAc mAbs and their application to the O-GlcNAc-modified proteome analysis of cells and tissues. These mAbs are efficient and specific towards O-GlcNAc, but not other glycosylation states. This strategy is simple and sensitive, allowing the rapid enrichment of large numbers of O-GlcNAc-modified peptides from complex biological samples. We believe these mAbs will provide a significant technical advancement in our ability to analyze and understand O-GlcNAc signaling.

## Abbreviations

GlcNAc: N-acetylglucosamine
GalNAc: N-acetylgalactosamine
HexNAc: N-acetylhexosamine
HexHexNAc: Hexose-N-acetylhexosamine
SA: sialic acid
CE: collision energy
ETD: electron transfer dissociation
HCD: higher-energy collisional dissociation
EThcD: electron transfer dissociation with supplemental HCD activation
pd: product-dependent
pd-EThcD: product dependent triggered EThcD
CID: collilsion induced dissociation
Abs: antibodies
WGA: wheat germ agglutinin
LC-MS/MS: liquid chromatography tandem mass spectrometry
mESCs: mouse embryonic stem cells
bRP: basic reversed-phase
XICs: extracted ion chromatograms

## Acknowledgments

We thank Matthew Fry for initial supplies of the antibodies, Namrata Udeshi and Susan Klaeger for technical assistance, Namrata Udeshi and Jason Maynard for critical assessment of the manuscript, and D.R. Mani, Karl Clauser and B. Hamilton for useful discussions. This work was supported by the Yale Science Development Fund (S.A.M., Yale) and by a postdoctoral fellowship from the California Institute for Regenerative Medicine (CIRM) Interdisciplinary Stem Cell Training Program II (to X.L.). This work was supported in part by grants from the National Cancer Institute (NCI) Clinical Proteomic Tumor Analysis Consortium grants NIH/NCI U24-CA210986 and NIH/NCI U01 CA214125 (to S.A.C.), and in part by NIH grant R35 CA210043 (to A.R.).

## Data availability

The original mass spectra may be downloaded from MassIVE (http:\\massive.ucsd.edu), MSV000087290, password Burt_2021.

---

**Supplemental Table 1** Byos/Byonic output for HexNAc-modified PSMs from data acquired from antibody-based enrichments of 1 milligram of mESC whole cell lysate using HCD-pd-EThcD.

**Supplemental Table 2** Byos/Byonic output for HexNAc-modified PSMs from data acquired from antibody-based enrichments of the synaptosomes using HCD-pd-EThcD.

**Supplemental Table 3** Byos/Byonic output for HexNAc-modified PSMs from data acquired from antibody-based enrichments of the synaptosomes using EThcD alone.

